# Alignment with sex-typical brain architecture: Links to mood pathology, sex hormones and transcriptomics of neurodegenerative conditions

**DOI:** 10.64898/2026.09.27.754773

**Authors:** Raluca Petrican, Alex Fornito

**Affiliations:** Institute of Population Health, Department of Psychology, University of Liverpool, Bedford Street South, Liverpool, L69 7ZA, UK; The Turner Institute for Brain and Mental Health, School of Psychological Sciences and Monash Biomedical Imaging, Monash University, Melbourne, VIC, Australia

**Keywords:** major depressive disorder, anxiety, ADHD, sex differences, sex hormones, neurodegeneration

## Abstract

Understanding the mechanisms that underpin the pervasive sex bias in risk for specific psychiatric and neurodegenerative conditions could substantially advance efforts promoting personalized medicine. As a first step towards addressing this issue, we applied imaging transcriptomics and multivariate analysis techniques to data from an adolescent sample enriched in anxiety/depression cases, a child sample enriched in attention deficit hyperactivity disorder (ADHD) cases, and adult patients with treatment resistant major depressive disorder (MDD). We tested whether, across sexes, alignment with female-typical, rather than male-typical, patterns of neural functional connectivity (FC) would relate to a diagnosis (adolescent sample) or severity of symptoms (adult sample) associated with internalizing disorders, which tend to be more often diagnosed in females. To provide preliminary evidence on its functional specificity, we compared the sex-aligned FC profile that distinguished neurotypical adolescents from those diagnosed with depression/anxiety against the sex-aligned FC patterns that distinguished neurotypical children from those diagnosed with ADHD. All the analyses controlled for age within each sample and sex.

Finally, we examined the spatial overlap between the sex-aligned FC profile linked to an internalizing diagnosis in adolescence and the transcriptomic maps of genes that show sex-dependent dysregulation in neurodegenerative conditions. Across sexes, increasing alignment with female-typical FC patterns, from sensory-perceptual to default mode regions, discriminated between typically developing and anxious/depressed teens and tracked longitudinal fluctuations in depressive symptoms within adult MDD patients. The identified internalizing-related gradient showed a negative correlation with risk for ADHD in the child sample. The brain regions most strongly aligned with female-, rather than male-, typical FC in depression/anxiety overlapped the spatial expression profiles of genes linked to gonadal hormone synthesis, as well as genes dysregulated in female autism spectrum disorder (ASD) and male neurodegenerative conditions, specifically, Alzheimer’s Disease (AD), Huntington’s Disease (HD) and Multiple Sclerosis (MS). The brain areas most strongly aligned with male-typical, rather than female-typical, FC patterns overlapped the transcriptomic maps of genes related to androgen and estrogen receptors, as well as of genes dysregulated in male ASD and female neurodegenerative conditions (i.e., AD, HD, MS). Our results raise the possibility that the link between mood pathology and neurodegeneration could reflect a partial, regional reversal of sex-normative functional brain profiles.

---

It is well-established that males have a higher risk for externalizing-spectrum disorders ^1,2^, while females have a higher risk for internalizing-spectrum disorders ^3^. Sex differences in psychiatric risk appear as early as childhood ^2^, but become established in early to mid-puberty^4,5^ when gonadal hormones play a reportedly key role in shaping the maturation of brain regions that are sexually dimorphic across species and implicated in the pathophysiology of disorders with sex-biased prevalence^5–7^. Understanding the mechanisms that drive sex differences in the prevalence of distinct psychiatric syndromes could substantially advance efforts to personalize detection and intervention paradigms.

Sex-dependent susceptibility to psychiatric disorders has traditionally been assumed to arise from categorical differences between males and females. However, this conceptual framework has been challenged by recent findings showing that sex differences in internalizing (i.e., anxiety) and externalizing (i.e., oppositional defiant disorder [ODD]) spectrum disorders are better explained by regional, rather than holistic, variability in relative alignment with female- vs male-typical FC^8^. Specifically, these studies have linked more severe anxiety symptoms, primarily among girls, to stronger alignment with female-typical FC in transmodal brain regions, whereas stronger alignment with male-typical FC in the same areas correlated with more severe ODD symptoms, especially among boys^8^.

This work has focused on the psychiatric implications of relative alignment with female-typical, rather than male-typical, FC among girls and relative alignment with male-typical, rather than female-typical, FC among boys. An important unresolved question is whether *differential* vulnerability to internalizing vs externalizing disorders between the sexes relates to the same regional patterns of alignment with female- vs male-typical FC. Two additional related issues remain unclear. One is whether longitudinal fluctuations in alignment with sex-typical functional brain architecture relate to longitudinal fluctuations in psychiatric symptoms. A robust association between the two would imply that they could be mechanistically related. The second question is whether patterns of alignment with sex-typical FC linked to psychiatric risk in early life could be meaningfully connected to sex-dependent neuropathology in older adulthood^9,10^, knowledge which could support the design of interventions that alleviate the impact of child mental ill-health on lifelong neurocognitive aging.

We addressed these issues by analyzing multimodal data from two child/adolescent samples enriched in internalizing (i.e., depression/anxiety) and externalizing (i.e., ADHD) cases, as well as a large group of adult patients with treatment resistant major depressive disorder (MDD). First, we characterized a profile of alignment with sex-typical functional connectivity (FC) patterns that distinguished neurotypical youths and those with an internalizing diagnosis, which we refer to as the Internalizing Neural Sexual Differentiation (iNSD) gradient. Given the female-biased prevalence of internalizing disorders, we expected that the iNSD gradient would primarily capture preferential alignment with female-, rather than male-typical FC patterns. This would be compatible with the interpretation that the greater prevalence of mood problems among female individuals may be partly due to the fact that these psychiatric symptoms are connected to functional brain features more often observed among female, rather than male, individuals. We focused on an internalizing-, rather than externalizing-, relevant gradient because mood disorders have been linked more robustly to sex differences in neurobiological circuits ^3,5,11^ relative to externalizing spectrum disorders such as ADHD ^12^.

Second, we explored the extent to which the iNSD gradient captures architectural brain features relevant to depression/anxiety, more specifically, as opposed to merely distinguishing between neurodivergent and neurotypical individuals. To this end, we examined the spatial overlap between the iNSD and patterns of alignment with sex-typical FC which distinguish between ADHD and normative development. Following the same reasoning as for the iNSD, given the male-biased prevalence of ADHD ^2^, we expected that children with ADHD (relative to neurotypical children) would show an overall pattern of stronger alignment with male-, rather than female-, typical FC pattern. Depending on whether distinct or overlapping brain regions contribute to the ADHD-related neural profile and iNSD, the two could be orthogonal or (somewhat) negatively correlated.

Third, we investigated whether longitudinal variability in the iNSD gradient tracked with the post-treatment persistence of depressive symptoms among MDD patients. Fourth, drawing on evidence that gonadal hormones are implicated in the emergence of sexually dimorphic brain structures^13,14^, we examined whether the iNSD gradient spatially correlated with the expression maps of genes regulating sex steroid synthesis and their corresponding receptors. Furthermore, given the well-documented involvement of sex hormones in typical and pathological neurocognitive aging (e.g.,^15–17^), we tested the potential relevance of the iNSD gradient to understanding the link between earlier life psychiatric conditions with sex-biased prevalence and neurodegenerative conditions which occur later in life and tend to be more often diagnosed in one sex. Leveraging the disorder-specific transcriptomic comparisons made available by refs.^18,19^, we focused on the sex-dependent transcriptomic dysregulation brain profiles observed in neurodegenerative conditions with sex-biased prevalence and pathophysiology, such as Alzheimer’s Disease (AD^16,20,21^ and multiple sclerosis (MS)^22,23^. We also investigated disorders showing sex differences in the underlying pathophysiology, but similar prevalence rates across males and females, such as Huntington’s Disease (HD)^24,25^ and frontotemporal dementia (FTD)^26^. These analyses aimed to determine whether regional patterns of relative alignment with female- vs male-typical FC, which are linked to the same internalizing symptoms across sexes in early life, would show distinct associations with the sex-dependent transcriptomic maps relevant to neurodegenerative risk.

## Methods and Materials

### Cohorts

We analysed data from three publicly available community samples (see Figure 2 for the selection criteria). One is an adolescent sample enriched in depression and anxiety cases (Boston Adolescent Neuroimaging of Depression and Anxiety [BANDA], N=176, aged 14-17 years, 60 boys). The second is a child sample enriched in ADHD cases (Oregon ADHD 1000, N =181, aged 7-12 years, 110 boys). The third is a sample of adult patients with treatment-resistant major depressive disorder (MDD) from the Perturbation of the Depression Connectome (PDC) study (N = 98, aged 20-76 years, 54 women) who underwent electroconvulsive therapy (ECT, N= 21), ketamine treatment (KET, N= 39) or overnight sleep deprivation (TSD, N = 38). Pre- and post-treatment symptom severity was assessed with the Hamilton Depression Rating Scale (HDRS^27^).

To minimize the confounding effect of motion^28^, we selected participants with relatively low in-scanner motion (i.e., average relative root mean square displacement < .20 mm). We further verified that the resting state data from all three samples, which had been processed with a version of the Human Connectome Project pipelines^29^, showed minimal evidence of contamination from motion artifacts (see Figure S10, cf^30^).

### Sex-Related FC Gradient

Sex-typical FC patterns were estimated for each parcel in a widely used functional atlas^31^. The reference group comprised healthy young adult participants (N = 336, 159 women, aged 22-30 years) in the Human Connectome Project (HCP), which were expected to show maximal differentiation of brain function based on their biological sex^8^. In the HCP, we regressed each of the 44850 FC estimates between each distinct pair of 300 brain regions against sex and a number of variables that had been linked to individual differences in FC and on which perfect matching between sexes is not feasible, specifically, age, income, employment status, race, handedness, average in-scanner motion, intelligence and psychopathology. The *t*-statistic corresponding to “sex” gauged alignment with sex-typical FC patterns because it quantifies how robustly a parcel-to-parcel FC index differentiates between men and women. Use of the *t*-statistic, rather than the corresponding beta value, safeguards against prioritizing parcels that show large beta values, but also large standard errors associated sex, suggestive of greater between-individual variability. Nonetheless, we verified that the sex-associated *t*-statistic and beta value corresponding to a given parcel-to-parcel FC index are very highly correlated (median *r* of .96, range: .79-.995). To confirm of the robustness of sex-related FC effects against the skewness of the confounds entered in the original regression analyses, we ran a Spearman’s partial correlation analysis between sex and each of 44850 FC estimates, while adjusting for all the nuisance variables entered in the corresponding regression analysis. As expected and compatible with the interpretation that data skewness was unlikely to bias the sex-relevant regression results, the correlation between the sex-related *t*-statistic from the original regression analysis and the sex-related Spearman’s rho for each of the 44850 FC estimates was very high (median *r* of .97, range: .85-.99). Note that the sex-related t-statistic values are used only for describing a sex-related FC strength gradient in the specific HCP sample pre-selected to act as a reference group because they are likely to show maximal differentiation in FC related to their biological sex. Because we are not making any inferences beyond the tested HCP sample, null hypothesis testing principles, such as thresholding based on *p*-values, are not applicable (Riffenburgh & Gillen, 2020).

### Alignment with the HCP Sex-Related FC Gradient

For each BANDA, Oregon or PDC participant, regional alignment with sex-typical FC patterns was computed as the rank-order correlation coefficient between each row of their own FC matrix (i.e., each parcel’s FC with each of the remaining parcels in the atlas) (Figure 1-F) and the corresponding row in the HCP *t*-statistic matrix (Figure 1-E). A positive coefficient indicates that an individual’s regional FC pattern better aligns with the ascending female-to-male FC strength pattern (suggestive of a more masculine FC strength profile); a negative coefficient implies alignment with the ascending male-to-female FC strength pattern (suggestive of a more feminine FC strength profile). We refer to the vector of such correlation values across regions as the individual’s neural sexual differentiation (NSD) profile. Because the HCP *t*-statistic matrix is based only on young adult FC profiles, both positive and negative rank order correlation coefficients reflect similarity to a young adult brain profile.

**Figure 1.**
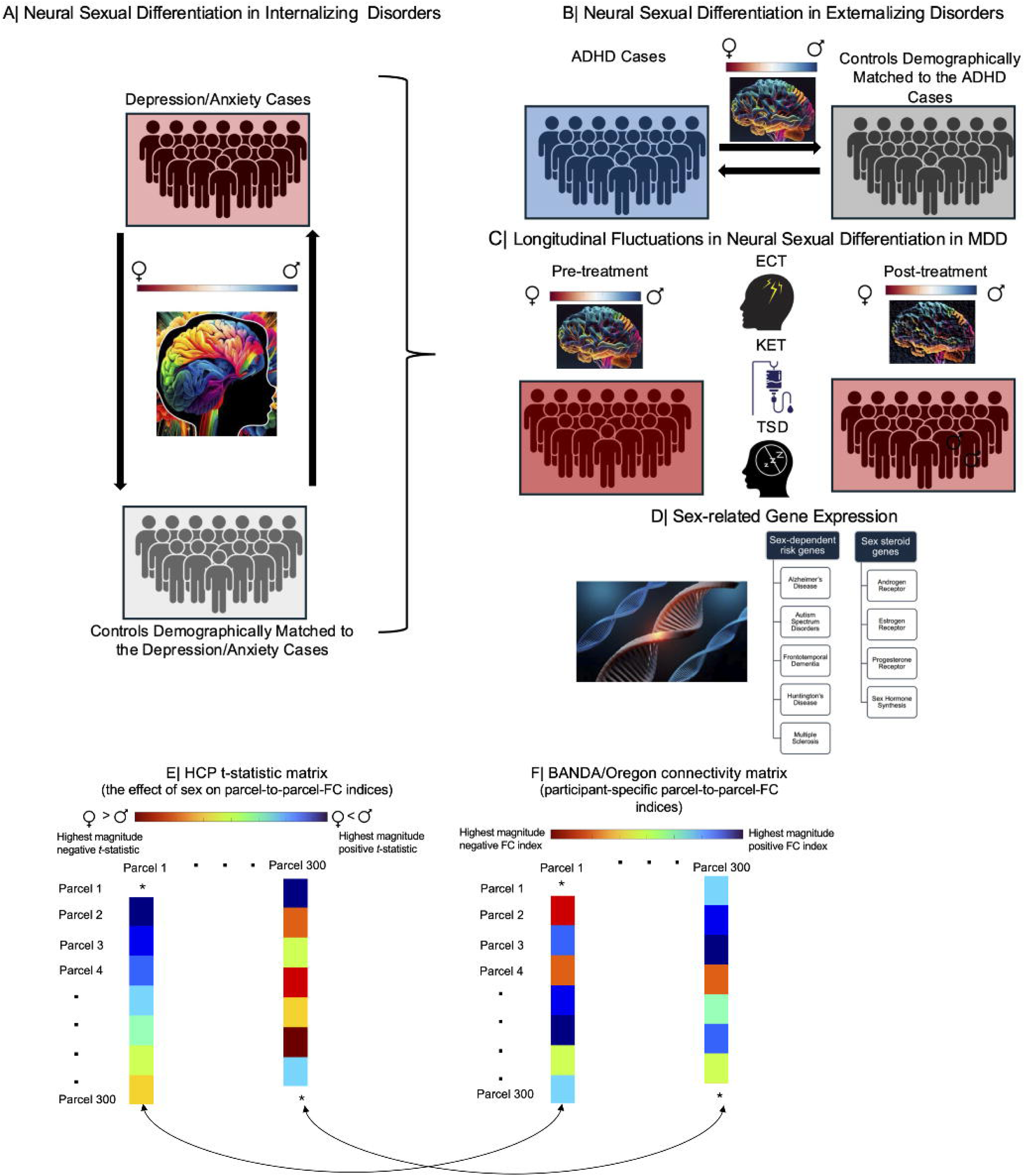
Overview of the Research Objectives. Our first objective was to identify regional patterns of alignment with sex-typical FC that distinguish neurotypical youths from those who had been diagnosed with an internalizing spectrum disorder (depression/anxiety), irrespective of their biological sex (panel A). As a second step, we examined the correlation between patterns of alignment with sex-typical FC connected to mood disorders (panel A) and those linked to a diagnosis of ADHD across both boys and girls (panel B). Third, we investigated whether the sex-related FC gradient in panel A could be used not only for cross-sectional comparisons, but also for longitudinal comparisons involving changes in symptom severity among MDD patients (panel C). Fourth, because gonadal hormones contribute to the emergence of sexually dimorphic brain structures and to vulnerability to neurodevelopmental and neurodegenerative conditions with sex-biased prevalence, we examined the correlation between the sex-related FC gradient in panel A and the transcriptomic maps of genes linked to sex steroid metabolism and risk for Alzheimer’s Disease, Autism Spectrum Disorder, Frontotemporal Dementia, Huntington’s Disease, and Multiple Sclerosis (panel D). Panels E-F depict the analytic steps through which alignment with female- vs male-typical FC patterns was estimated. *T*-statistic values corresponding to self-connections (panel E) or FC indices corresponding to self-connections (panel F) were set to zero. ADHD = attention deficit hyperactivity disorder. MDD = major depressive disorder.

**Figure 2.**
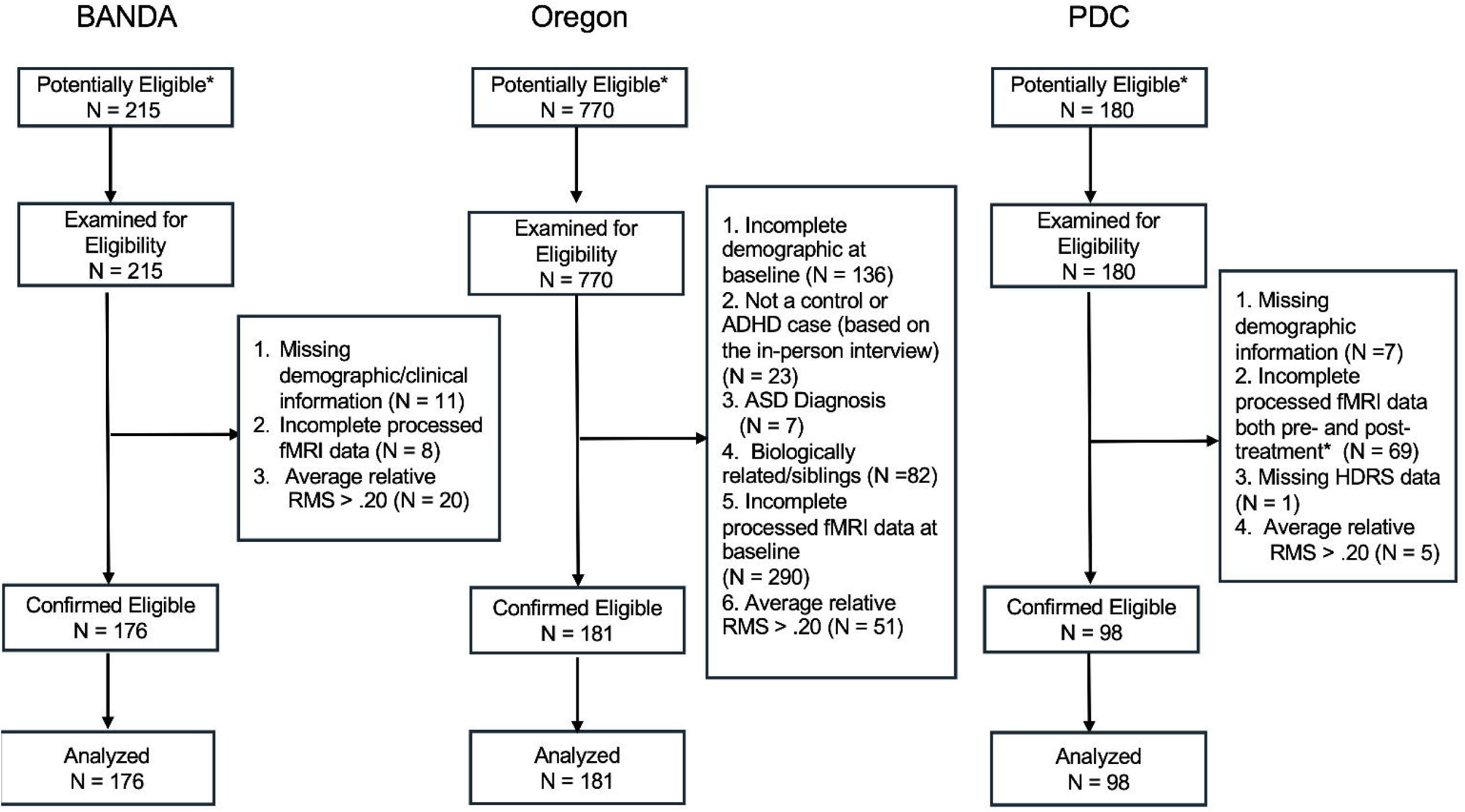
STROBE flowchart describing sample selection for the BANDA, Oregon ADHD 1000 and PDC cohorts.

Note that principles of null hypothesis testing (e.g., thresholding based on *p*-values) do not apply to these correlation analyses which merely gauge the similarity between two measured variables (i.e., corresponding columns in the HCP t-statistic matrix and each participant’s FC matrix).

We replicated all the results with HCP sex-related FC models further adjusted for brain size and individual differences in female hormonal state. In supplemental analyses, we confirmed that both sets of HCP sex-related FC models can robustly distinguish between sexes in the adolescent, child and adult samples. Specifically, in all three samples, the male participants showed stronger alignment in FC with the HCP men, rather than the HCP women, whereas the female participants showed stronger alignment in FC with the HCP women, rather than the HCP men (see Supplemental Methods and Figures S11-12).

### Gene Expression Profiles

We leveraged the microarray data released by the Allen Brain Institute and estimated in abagen^32^ brain-wide expression of the 25 steroid genes identified by ref.^14^ and of the 11, 976 genes showing sex-dependent transcriptomic dysregulation in the brain cells of AD, FTD, HD, MS, or ASD patients (relative to matched controls)^19^. The disorder-related genes are part of ZEBRA^19^, a transcriptomic atlas that integrates gene expression data from 33 studies, including over 4 million human and murine cells from 39 brain regions. We included only the ZEBRA genes which have been shown by ref.^18^ (Supplementary Dataset 4) to exhibit statistically robust transcriptomic dysregulation in neuronal and non-neuronal (e.g., glial, vascular) brain cells as a function of sex and disorder. The gene expression output from abagen was parcellated based on the functional atlas used in our neuroimaging analyses. For the sake of simplicity, given the availability of microarray data from all six Allen Brain Institute donors, we only used left hemisphere data.

To ease interpretation, we aggregated the expression levels of the sex hormone-related genes based on four broad categories: three related to receptor types (androgen, estrogen, progesterone receptor family) and one related to sex steroid synthesis (see Table S1 for the gene list and chromosome locations). The normative transcriptomic maps of genes showing differential expression as a function of disorder (AD, ASD, FTD, HD, MS) and sex of the patient were estimated by averaging the corresponding expression levels within each parcel, separately by disorder and sex. This resulted in two matrices: one contained the normative spatial expression of genes that are dysregulated in female patients with AD, ASD, FTD, HD, and MS relative to controls, whereas the second encompassed the normative spatial expression of genes dysregulated in male patients with AD, ASD, FTD, HD, and MS relative to controls. Subsequently, we subtracted the female matrix from the male matrix.

This yielded a matrix of 5 (disorders) x 146 (parcels), which comprised the sex-differential normative expression maps for genes that are dysregulated in AD, ASD, FTD, HD, and MS. In this matrix, positive values indicate that the respective parcel shows relatively greater normative expression of genes that are dysregulated in male (rather than female) patients for one of the five disorders, whereas negative values indicate that the respective parcel shows greater normative expression of genes that are dysregulated in female, rather than male, patients.

## Statistical Analysis

We used two multivariate data-driven techniques, specifically, canonical correlation analysis (CCA) and partial least squares correlation analysis (thereafter, PLS), which explain commonalities in two sets of variables by creating linear combinations, dubbed “variates” (CCA) or “latent variables” (LVs, PLS), from the observed variables in each set ^33^. Our decision to rely on PLS and CCA was grounded in evidence that psychiatric conditions are best characterized in relation to intrinsically correlated functional systems spanning multiple anatomical brain regions ^34–37^.

All the multivariate models described below were implemented with 10-fold cross-validation (i.e., the model discovery and testing were performed on non-overlapping subsets of the same sample). Statistical significance was established through permutation testing (100,000 samples), whereas the reliability of the model estimates (99% confidence intervals [CIs] for within-set relationships and 95% CI for between-set relationships) was quantified via bootstrapping (100,000 samples).

The iNSD gradient was intended to capture the co-occurring brain-wide patterns of regional alignment with female- vs male-typical FC which distinguish clinical depression/anxiety from normative development. Consequently, this gradient was estimated with PLS because it outperforms CCA on reliability and reproducibility in datasets containing a large number of highly correlated within-set variables ^33,38^. To test the overlap between the iNSD gradient and individual male- vs female-typical FC patterns linked to ADHD and MDD symptom persistence across sexes, we relied on partial correlations because they provide a more direct estimate of brain-behaviour associations relative to PLS which maximises within- set correlations ^33^. We relied on Spearman’s correlations because of the marked skewness of the HDRS variables in the PDC sample (for the analyses involving the dummy-coded ADHD diagnosis variable, Spearman’s correlations constitute the default option). CCA was not a viable option for computing these brain maps because of the number of neural variables involved ^39^. CCA was though recruited to quantify the overlap between the iNSD gradient and the brain maps tracking longitudinal variations in MDD symptom severity, as well as the spatial expression maps of genes relevant to neurodegeneration and sex steroid function. As specified above, CCA was implemented with a 10-fold cross-validation procedure. Hence, if the skewness of any of the scrutinized variables had biased the extracted CCA mode (i.e., the associated standardized coefficients for the identified variates) in any of the training folds, the cross-validation tests would have failed.

## Results

### Regional alignment with sex-typical FC patterns relates to internalizing disorders

We started by examining whether patterns of NSD can discriminate between neurotypical youths and youths diagnosed with an internalizing disorder in the BANDA sample. To this end, we conducted a PLS analysis in which we related each adolescent’s parcel-specific alignment with the HCP sex-related FC gradient to their corresponding diagnostic group, either depression/anxiety, N = 121, 35 boys, or controls, N = 55, 25 boys.

The discovery PLS analysis conducted on raw/non-residualized data across all training folds extracted one diagnostic group-brain latent variable (LV) pair (*r*_disc_ of .38, permutation-based *p* = .010, 95% confidence interval [CI] = [.32; .58]), which was successfully cross-validated (*r*_CV_ of .17, permutation-based *p* = .024, 95% confidence interval [CI] = [.02;.32]), after controlling for sex, age, in-scanner motion, parental education, handedness and racial background. This LV constituted an internalizing-related NSD gradient (i.e., iNSD) in which depressed/anxious adolescents, regardless of sex, show increasingly stronger alignment with female-, rather than male-, typical FC from sensory/perceptual (i.e., visual, somatomotor) to association (i.e., default mode network [DMN], Control) regions (Figure 3-B, C, E, F).

**Figure 3.**
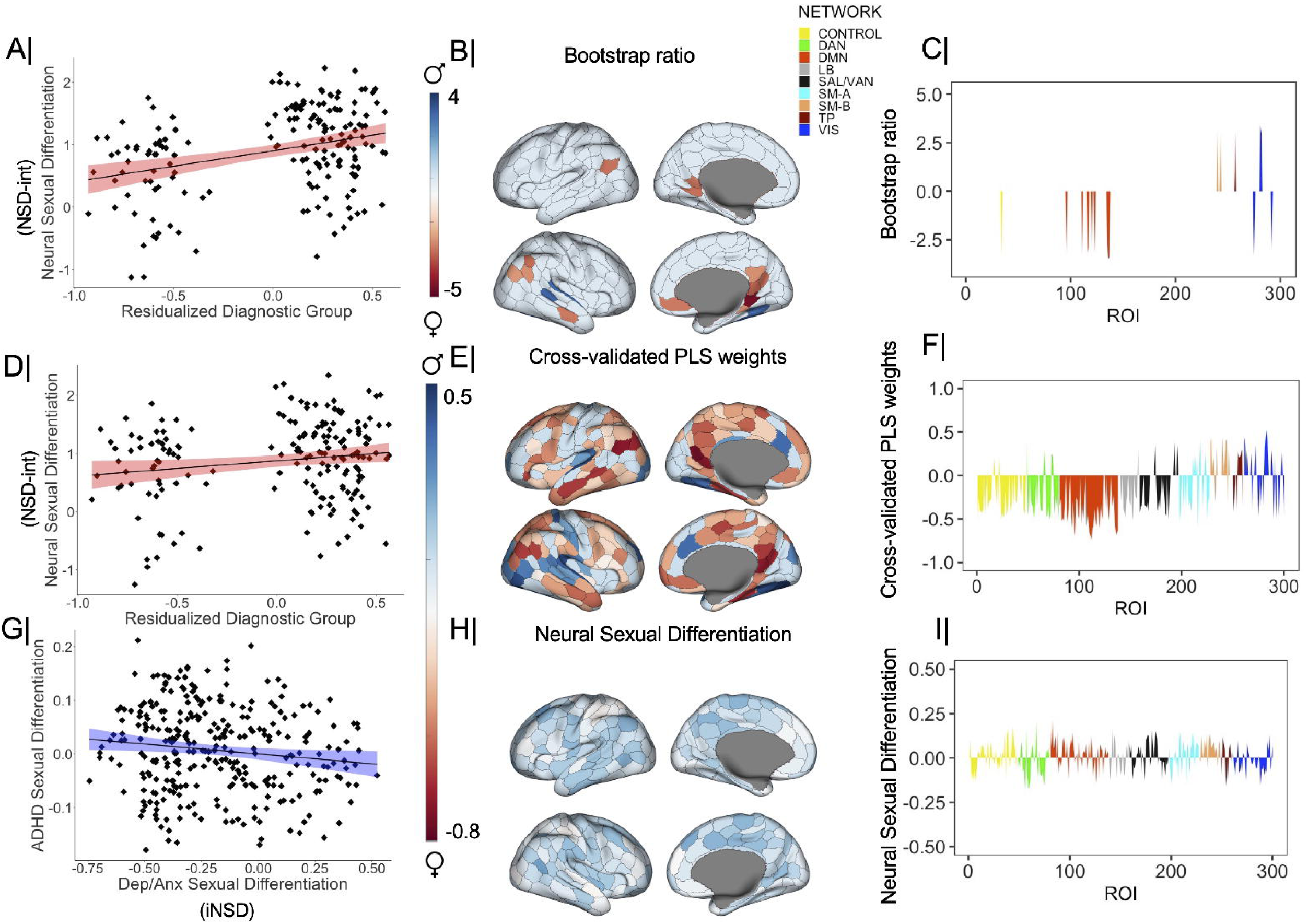
Patterns of alignment with sex-typical FC linked to an internalizing vs externalizing diagnosis. Panels A and D show the correlation between the diagnostic group variable (coded 0 for controls, 1 for adolescents diagnosed with depression or anxiety), which had been adjusted for the confounders listed under “Control variables”, and the brain LV from the discovery PLS (panel A) or the cross-validation procedure (panel D). Panel B depicts the ROI-specific weights/loadings on the brain LV identified with the discovery PLS analysis with a bootstrap ratio greater than 2.75 in absolute value (equivalent to a 99% CI). Panel C presents the same information aggregated by the Schaefer atlas networks. Panel E depicts the Schaefer ROIs robustly correlated (based on cross-validated 99% CIs) with the predicted value of the brain LV from the cross-validation procedure. These are partial correlations controlling for the confounders listed under “Control variables”. Panel F presents Schaefer network-based distributions of partial correlations summarizing the ROI-specific results from panel (E). Panel G shows the correlation between the cross-validated and unthresholded brain LV from the PLS analysis depicted in panels A-F (dubbed ‘iNSD” in the main text) and the map of alignment with sex-typical FC related to ADHD (adjusted for the confounders listed under “Control variables”). Panel H shows the correlation between ADHD (coded 1 for cases, 0 for controls) and the alignment of each Schaefer parcel with sex-typical FC patterns. Panel I depicts the same information as panel H but aggregated by the Schaefer networks. The subnetworks from the Schaefer 17-network atlas (e.g., Control A/B/C) have been combined into one to increase comparability with the Gordon atlas. In panels B, C, E, F, H, and I positive values indicate stronger alignment with male-, rather than female-, typical FC, whereas negative values indicate stronger alignment with female-, rather than male-, typical FC for a specific parcel. As stated in the main text, alignment with sex-typical FC patterns was estimated in reference to resting state connectivity data from the Human Connectome Project. BSR = bootstrap ratio. PLS = partial least squares. LV= latent variable. CI = confidence interval. Schaefer networks: TP= temporo-parietal. SAL-VAN = salience/ventral attention. LB = limbic. DMN = default mode. DAN = dorsal attention. SM-A = somatomotor-A. SM-B =somatomotor-B. VIS = visual.

Across sexes, relative alignment with female- vs male-typical FC patterns is reportedly related to adrenarche but not gonadarche^8^. To probe this relationship, we tested the correlation between scores on the identified iNSD gradient and the youths’ Tanner developmental stage scores (adjusted for the same nuisance variables entered in the PLS analysis above). There was no robust association between the two, *r* = -.14, 95% CI = [-.30; .04], permutation-based *p* = .08, which implies that the iNSD gradient does not merely capture individual differences in pubertal timing.

### Alignment with male-typical FC patterns distinguishes children with ADHD from neurotypical children

Next, we explored whether the iNSD gradient captures FC patterns specifically related to internalizing disorders, rather than broad neurodivergence. Consequently, we examined the relationship between the iNSD gradient and patterns of alignment with sex-typical FC that distinguish children with ADHD from neurotypicals. Spearman’s partial correlations quantified the relationship between diagnostic group (ADHD, N = 107, 74 boys, and controls, N = 74, 36 boys) and the alignment of each Schaefer parcel with the HCP female-to-male FC gradient, after adjusting for sex, age, parental education, racial background and in-scanner motion.

A robust, yet modest, negative correlation emerged between the cross-validated iNSD LV and the ADHD partial correlation map, *r* = -.13, permutation-based *p* = .048, 95% confidence interval [CI] = [-.03; -.23]. Inspection of Figure 3-H, I suggested that this relationship is driven by transmodal DMN regions which show a more male-typical FC profile in ADHD and a more female-typical FC profile in internalizing disorders, compared to normative development.

### Longitudinal increases in iNSD track fluctuations in depression symptoms

Having shown that relative alignment with female- vs male-typical FC patterns relates to the onset of depression/anxiety across sexes, and to a lesser extent, ADHD, we next tested whether longitudinal changes in alignment with sex-typical FC patterns track the post-treatment persistence of mood symptoms in a clinical cohort. Our analyses focused on 98 MDD patients from the PDC study, before and after they underwent one of three treatments: electroconvulsive therapy (ECT, N = 21, 13 women), ketamine infusion therapy (KET, N = 39, 19 women) or 24-hour sleep deprivation (TSD, N = 38, 22 women). For each patient, longitudinal changes in regional patterns of alignment with sex-typical FC were computed by subtracting the pre-treatment standardized score of each parcel from the corresponding post-treatment standardized score. Symptom difference scores on each HDRS subscale were similarly computed. Spearman’s partial correlation analyses, adjusted for treatment type, age at treatment, pre-to-post-treatment delay, sex, race, handedness and average in-scanner motion, quantified the relationship between parcel-specific changes in alignment with the HCP female-to-male FC gradient and changes in the HDRS subscale scores from pre- to post-treatment. In the resulting correlation matrix, a positive coefficient indicates that longitudinal increases in depressive symptom severity are linked to longitudinal increases in alignment with male-, rather than female-, typical FC, while a negative correlation coefficient suggests that pre- to post-treatment worsening of MDD symptoms relates to increasing alignment with female-, rather than male-, typical FC.

To determine whether the iNSD gradient overlaps with patterns of alignment with sex-typical FC which track persistence in depressive symptoms, we conducted a CCA linking the cross-validated brain LV from the PLS analysis (Figure 3-E) to the above described partial correlation maps.

The cross-validated CCA mode (*r*_CV_ = .72, 95% CI = [.67; .77], *p_spin_* = 10^-5^) indicated an overlap between the iNSD gradient and patterns of alignment with female-typical FC connected to post-treatment symptom persistence in the MDD sample. In other words, longitudinal increases in alignment with female-, rather than male-, typical FC, particularly within the DMN and Control systems (Figure 3-B, C, E, F), correlated with longitudinal increases in depressed mood, anxiety (psychic, somatic), lethargy (i.e., low somatic energy), agitation, guilt, suicidal tendencies and anhedonia (i.e., reduced interest in work and personal activities) among the MDD patients (Figure 4-A). Widespread longitudinal increases in alignment with male-, rather than female-, typical FC were associated with worsening of appetite-related problems.

**Figure 4.**
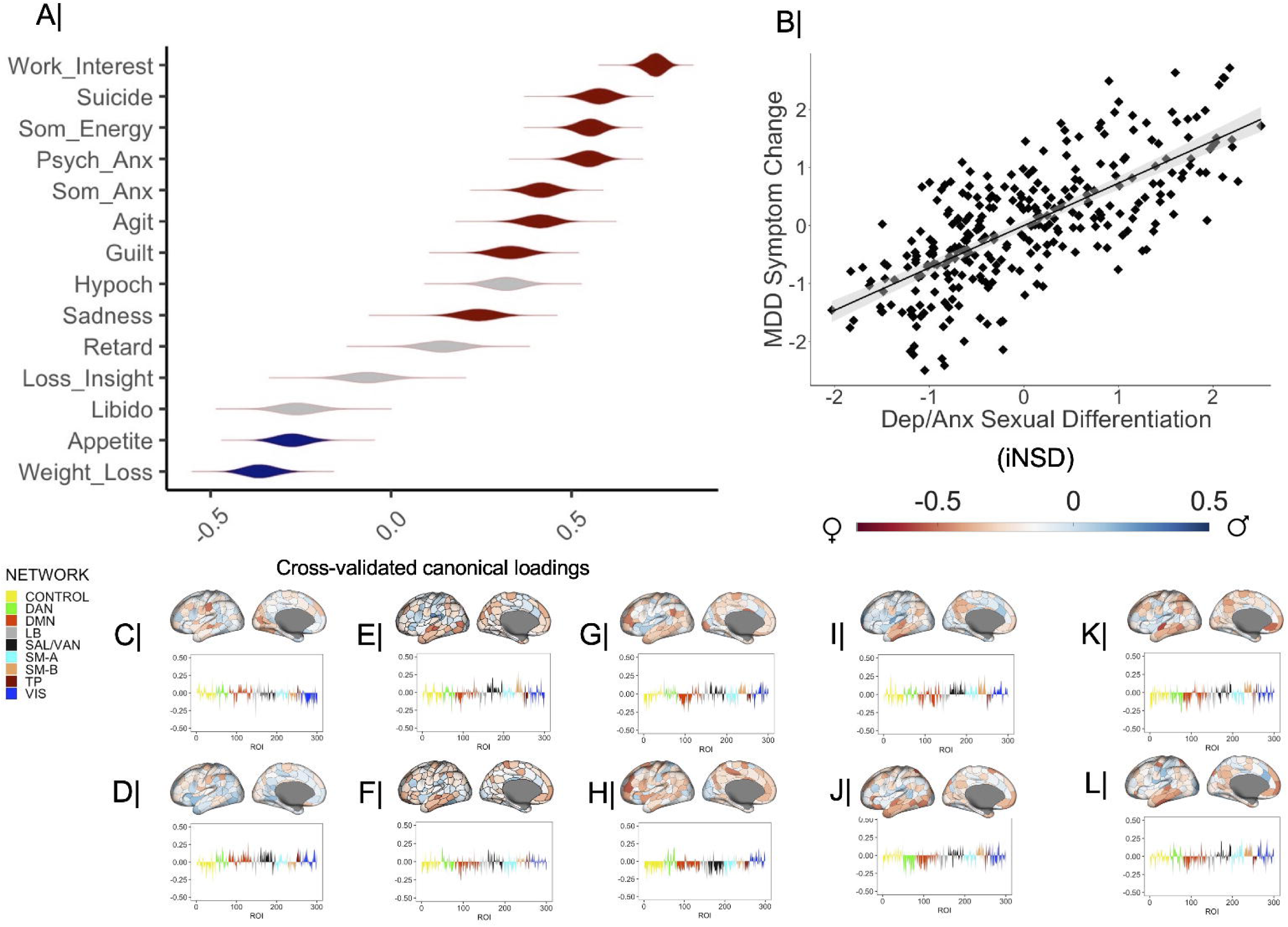
Longitudinal fluctuations in alignment with sex-typical FC patterns and MDD symptoms. Panel A shows the correlation coefficients describing the relationship between alignment with sex-typical FC connected to longitudinal changes on the HDRS subscales and the predicted value of their corresponding canonical variate across all test CCA folds. The violin plots in panel A depict the distribution of correlation coefficients across the 100,000 bootstrap samples from the cross-validation procedure. Gray colored violin plots indicate non-robust canonical loadings (based on the bootstrapped 99% CIs from the cross-validation procedure). Panel B contains the scatter plot describing the linear relationship between the cross-validated, unthresholded iNSD gradient and the CCA variate pictured in panel A. Panels C-L depict the correlation maps between pre- to post-treatment changes in alignment with sex-typical FC for each Schaefer parcel (top) or network (bottom), and changes in scores on the HDRS subscale, specifically, weight loss (C), appetite (D), sadness (E), guilt (F), agitation (G), somatic anxiety (H), psychic anxiety (I), somatic energy (J), suicidality (K) and work/personal interests (L). CCA = canonical correlation analysis. CI = confidence interval. HDRS = Hamilton Depression Rating Scale. LV = latent variate. PLS = partial least squares correlation analysis. Anx = anxiety. Dep = depression. Hypoch = hypochondriasis. Psych = psychic. Retard = retardation (i.e., slowness). Som = somatic. Schaefer networks: TP= temporo-parietal. SAL-VAN = salience/ventral attention. LB = limbic. DMN = default mode. DAN = dorsal attention. SM-A = somatomotor-A. SM-B =somatomotor-B. VIS = visual.

## Transcriptional correlates of alignment with sex-typical FC

Sex differences in brain structure are accentuated in areas characterized by high expression of sex steroid genes ^14^. We therefore tested whether our iNSD gradient would align with the spatial expression maps of sex steroid genes, such that areas that show the strongest alignment with sex-typical FC would also show the strongest expression of sex steroid genes from one or more of the four scrutinized categories. Gonadal hormones reportedly contribute to sex-differential trajectories of both normative and pathological aging (e.g.,^15–17^). Therefore, we were also interested in whether the iNSD gradient would overlap with the normative spatial expression maps of genes that show sex-differential dysregulation in neurodegenerative conditions (i.e., AD, FTD, HD, MS).

A CCA linked the iNSD (Figure 3-E) to the four normative spatial expression maps of the sex steroid genes and the five normative expression difference maps for genes dysregulated among male vs female patients with AD, FTD, HD, MS, ASD. The cross-validated mode emerging from this analysis (*r*_CV_ = .25, 95% CI = [.07; .40], permutation-based *p_spin_* = .008) (Figure 5-B) indicated that the brain areas most strongly aligned with female-typical FC in depression/anxiety (mostly, DMN and Control), tend to show greater normative expression of genes modulating sex steroid synthesis, as well as relatively stronger normative expression of genes that are dysregulated in male neurodegeneration rather than those that are dysregulated in female neurodegeneration^18,19^ (Figure 5-A). In contrast, sensory regions which show relatively greater alignment with male-, rather than female-, typical FC in depression/anxiety are characterized by higher normative expression of androgen and estrogen receptor genes, as well as of genes dysregulated in female, rather than male, neurodegeneration (Figure 5-A). With regards to ASD, the CCA output revealed that greater normative expression of genes dysregulated in male ASD typifies the brain regions with the strongest male-aligned FC, whereas greater normative expression of genes dysregulated in female ASD was observed in the brain regions with the strongest female-aligned FC.

**Figure 5.**
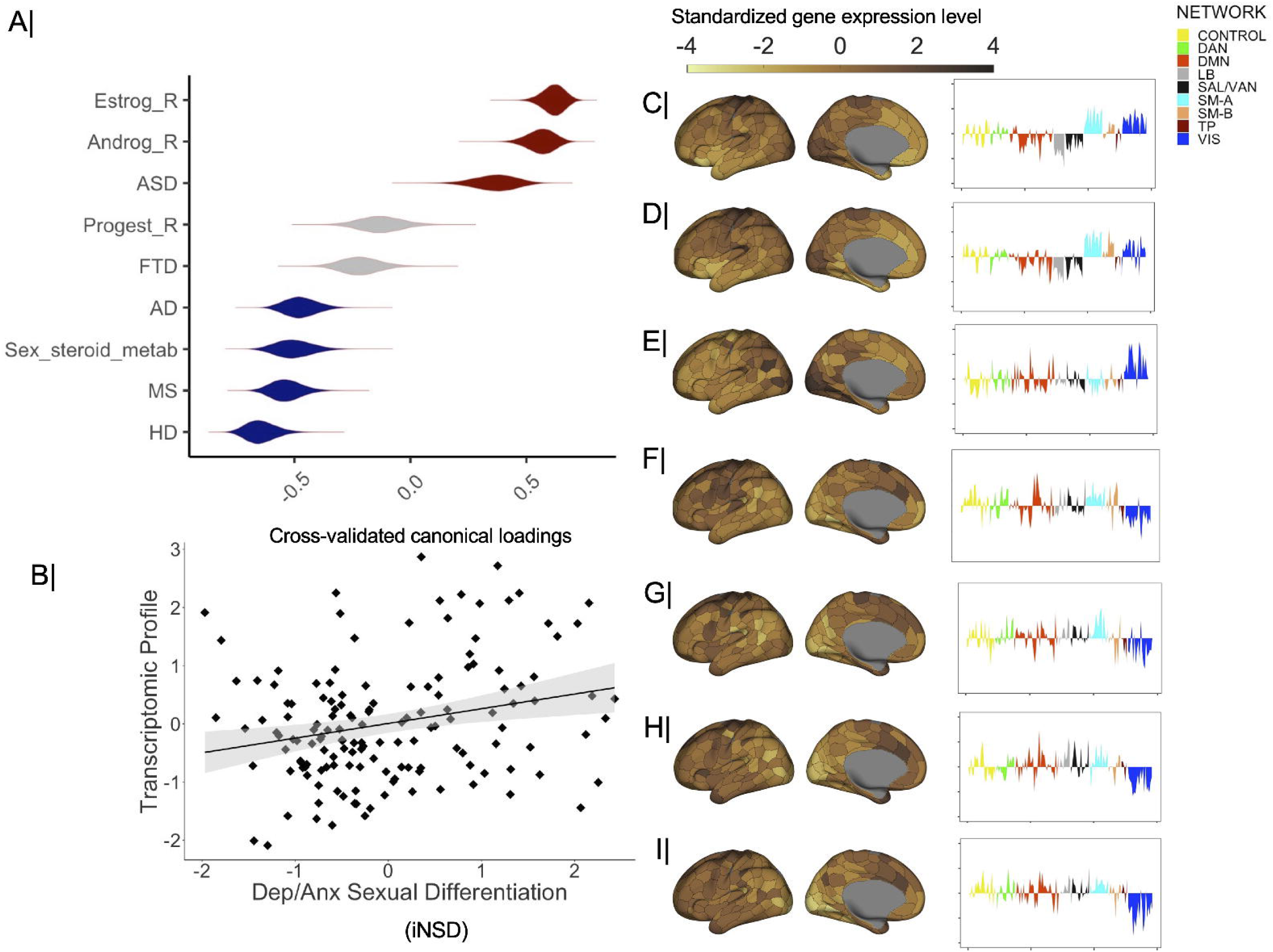
Transcriptomic correlates of alignment with sex-typical FC patterns linked to an internalizing diagnosis. Panel A shows the correlation coefficients describing the relationship between the sex steroid- and disorder-relevant transcriptomic maps and the predicted value of their corresponding canonical variate across all test CCA folds. The violin plots in panel A depict the distribution of partial correlation coefficients across the 100,000 bootstrap samples from the cross-validation procedure. Gray colored violin plots indicate non-robust canonical loadings (based on the bootstrapped 99% CIs from the cross-validation procedure). Panel B contains the scatter plot describing the linear relationship between the cross-validated, untresholded iNSD gradient and the CCA variate pictured in panel A. Panels C-I depict the parcel- and network-specific standardized expression values for estrogen receptor genes (A), androgen receptor gene (B), sex-differential ASD risk genes (C), progesterone receptor genes (D), sex-differential FTD risk genes (E), sex-differential AD risk genes (F), genes related to sex steroid synthesis/metabolism (G), sex-differential MS risk genes (H) and sex-differential HD risk genes (I). iNSD = neural sexual differentiation (related to) internalizing symptoms. LV = latent variable. CI = confidence interval. Metab = metabolism. _R = receptor. Androg = androgen. Estrog = estrogen. Progest = progesterone. AD = Alzheimer’s Disease. ASD = Autism Spectrum Disorder. FTD = Frontotemporal Dementia. HD = Huntington’s Disease. MS = Multiple Sclerosis. Schaefer networks: TP= temporo-parietal. SAL-VAN = salience/ventral attention. LB = limbic. DMN = default mode. DAN = dorsal attention. SM-A = somatomotor-A. SM-B =somatomotor-B. VIS = visual.

To test the specificity of the brain-transcriptomic profile association, we used the toolbox developed by ref.^40^ and the 15,632 brain-expressed genes released by the Allen Brain Institute to generate via random selection 100,000 gene sets containing the same number of genes as the sex steroid and disorder-specific (i.e., female AD, male AD, female ASD, male ASD, female FTD, male FTD, female HD, male HD, female MS, male MS) sets entered in the CCA above. Random generation of these additional gene sets was considered to be an appropriate strategy since across the 146 left hemisphere Schaefer ROIs with available transcriptomic data, the mean co-expression level in the original sex steroid gene set was .04, whereas in the sex-dependent and disorder-specific gene sets, it ranged from .01 to .03.

Using the same procedure as for the original gene sets, we estimated the normative expression maps for the four sex-steroid and five sex-differential disorder-specific transcriptomic aggregates across the 100,000 randomly generated sets. For each set, we computed transcriptomic profile scores by applying the standardized coefficients from the CCA conducted on the original gene sets (the coefficients were averaged across the 10 training folds). The correlation between the iNSD gradient and the transcriptomic profile thus computed was lower than the one observed with the original cross-validated transcriptomic profile for 99,992 of the 100,000 randomly generated gene sets, a result that broadly supports the specificity of our reported brain-gene associations.

## Discussion

Characterization of the sex-dependent mechanisms through which environmental and genetic factors shape psychiatric vulnerability constitutes a key step in personalizing clinical detection and intervention paradigms. Leveraging an adolescent multimodal dataset, we provided evidence that regional variability in relative alignment with female- vs male-typical FC patterns can discriminate between normative development and a clinical diagnosis of depression/anxiety regardless of one’s biological sex and independent of reproductive maturation stage. A modest, yet reliable negative association was observed between the internalizing-relevant brain gradient and the patterns of alignment with sex-typical FC distinguishing individuals with ADHD and neurotypicals. This effect, which provides suggestive evidence on the specificity of the iNSD-internalizing symptom relationship, appeared to be driven by complementary patterns of alignment with sex-typical FC in the DMN, such that increased alignment with female-typical FC was linked to depression/anxiety and increased alignment with male-typical FC was linked to ADHD.

In a third adult dataset, we demonstrated that longitudinal fluctuations along the internalizing-relevant gradient identified in the adolescent sample are coupled with longitudinal fluctuations in depressive symptoms among men and women diagnosed with MDD. Specifically, a diagnosis of depression/anxiety and post-treatment persistence of depressed mood, anxiety, agitation, lethargy, guilt, suicidal ideation and anhedonia were most robustly associated with female-, rather than male-, aligned FC in the DMN and Control networks, along with male-aligned FC in visual and somatomotor systems. The reverse pattern of sex-related alignment correlated with more severe appetite-related problems after MDD-related treatment. Our findings thus imply that regional alignment with sex-typical FC can provide useful insights into individual differences in psychiatric symptoms with sex-biased prevalence, including their longitudinal fluctuations.

Using microarray data from the Allen Human Brain Atlas, we demonstrated that the iNSD gradient aligns with the normative spatial expression maps of sex steroid genes and of genes that show sex-dependent dysregulation in neurodegenerative conditions. Specifically, the regions with the strongest female-aligned FC in depression/anxiety, which are mainly within the DMN and Control network (Figure 3-E), are characterized by greater expression of genes implicated in sex steroid synthesis and of genes that are dysregulated in male patients, rather than those dysregulated in female patients, with a neurodegenerative condition (AD, HD and MS). Moreover, visual and sensory regions, which show the strongest alignment with male-typical FC patterns in depression/anxiety, tend to be enriched in androgen and estrogen receptor-related genes, as well as in genes that are dysregulated in female (rather than male) patients with AD, HD or MS. Our findings resonate with existing literature on the contribution of gonadal hormones to internalizing risk^15,41^, vulnerability to pathological brain aging^16,17,42,43^, as well as evidence linking mood disorders to AD^9,44^, HD^24,45^ and MS^46,47^. The complementary neuroimaging transcriptomic patterns linking mood pathology to male vs female neurodegeneration fit well with the well-documented sex differences in the pathophysiology of AD^16,48–50^, HD^24,25,51^ and MS^52–55^.

Recall that brain regions most strongly aligned with female-typical FC in depression/anxiety showed greater normative expression of genes dysregulated in male (rather than female) neurodegeneration, whereas areas least strongly aligned with female-typical FC in depression/anxiety showed greater normative expression of genes dysregulated in female (rather than male) neurodegeneration. In light of cross-species evidence implicating gonadal hormone signalling in neural sexual differentiation^13^, our present findings raise the possibility that regional deviations from neurohormonal profiles typical of one’s biological sex may link mood pathology to vulnerability to later neurodegeneration. This argument is broadly compatible with existing evidence on the role of altered androgen signalling in AD-related deficits^42^, including the progression of cognitive and neuroimmune deficits in male AD^56^, as well as the role of estrogen availability and signalling in normative and pathological brain aging trajectories, such as AD, among women^16^. While our findings imply that the link between mood pathology and neurodegeneration may be related to regional alterations in sex-typical brain function, the relationship between depression/anxiety and ASD seems to be connected to regional deviations in the direction of sex-typical neural profiles. Specifically, stronger normative expression of genes dysregulated in male (rather than female) ASD was observed in the brain regions most strongly aligned with male-typical FC in depression/anxiety, whereas stronger normative expression of genes dysregulated in female (rather than male) ASD was primarily detected in the regions showing the strongest alignment with female-typical FC. These results raise the possibility that the well-documented comorbidity between ASD and mood disorders^57^, particularly during childhood and adolescence, may be linked to slight deviations in alignment with female- vs male-typical FC, which remain nonetheless concordant with the direction of maturational processes typical of one’s biological sex.

Our study has several limitations which can inform future investigations. First, the identification of the iNSD, showing broadly complementary relationships with ADHD and depression/anxiety, was based on a cross-sectional sample. Incorporation of longitudinal data would shed light on whether fluctuations in relative alignment with female- vs male-typical FC could be used to monitor the progression of both internalizing and externalizing spectrum disorders, including treatment-related improvements in symptoms. Second, the iNSD gradient was identified in an adolescent sample. Future studies using a wider age range for the discovery analyses would help in determining whether the patterns of alignment with sex-typical FC, which are connected to internalizing vs externalizing disorders, change across the life course as a function of experience and neurocognitive aging. Third, the size of the MDD patient sample precluded investigation of effects specific to each treatment type. Extant evidence points to distinguishable neurobiological pathways through which ECT, KET and TSD induce therapeutic change^58–63^. Future investigations with sufficiently large samples for each treatment type could provide key insights into the multiscale processes through which ECT, KET and TSD lead to yoked alterations in patterns of alignment with sex-typical FC and depressive symptoms. Fourth, our analyses relied on the Allen Human Brain Atlas for transcriptional correlates and, thus, we could not directly investigate individual differences. Studies probing the relationship between individual differences in the expression of these genes (e.g., via methylation patterns, cross-species investigation of epigenetic regulators) and ADHD/mood-relevant alignment with sex-typical brain architecture would represent an important first step towards identifying the multiscale mechanisms underpinning vulnerability to disorders with sex-biased prevalence.

In conclusion, stronger alignment with female-typical, rather than male-typical, FC patterns, particularly in transmodal brain regions, was associated with a clinical diagnosis of an internalizing disorder in both males and females. Longitudinal fluctuations in the degree of alignment with female-, rather than male-, typical FC covaried with fluctuations in adult MDD symptom severity. Spatial transcriptomic analyses implied that the identified iNSD gradient could reflect individual differences in sex steroid function, while also raising the possibility that regional deviance from sex-typical neurohormonal profiles could link earlier mood pathology to brain degeneration in later adulthood. Our results speak to the relevance of sex-dependent functional brain architecture for characterizing multiscale contributors to psychiatric conditions during maturation and their potential contribution to sex differential trajectories of neurocognitive aging.

## Materials & Correspondence

Correspondence and material requests should be addressed to R.P.

## Data availability

The raw data are available at https://nda.nih.gov/edit_collection.html?id=3037 (DOI: <u>10.15154/3tk5-pb47</u>) (BANDA) https://nda.nih.gov/study.html?id=1938 (DOI: 10.15154/1528485) (Oregon), https://nda.nih.gov/ccf/disease-studies (DOI: 10.15154/00b0-hw43) (PDC) and at https://db.humanconnectome.org (HCP) upon completion of the relevant data use agreements. **Code availability.** We used already existing code, as specified in the main text with links for free download.

## Acknowledgments

Data used in the preparation of this article were obtained from the Boston Adolescent Neuroimaging of Depression and Anxiety (BANDA) study, the Oregon ADHD-1000: A longitudinal data resource enriched for clinical cases and multiple levels of analysis, the Perturbation of the Depression Connectome (PDC) Study, and the Human Connectome Project, WU-Minn Consortium (Principal Investigators: David Van Essen and Kamil Ugurbil; 1U54MH091657; funders: the 16 NIH Institutes and Centers that support the NIH Blueprint for Neuroscience Research and the McDonnell Center for Systems Neuroscience at Washington University). The BANDA project was supported by the National Institute of Mental Health, U01MH108168 (Gabrieli, Whitfield-Gabrieli), and F32MH114525 (Hubbard). The project was made possible by the resources provided by Shared Instrumentation Grants 1S10RR023401, 1S10RR019307, and 1S10RR023043. The Oregon ADHD-1000 project was supported by multiple NIMH grants: 1R01-MH59105 (Nigg); R01-MH63146 (Nigg); R01-MH070004 (Friderici); 1R01MH86654 (Nigg-Fair); R01MH099064 (Nigg); 1R37MH59105 (Nigg); 2R56MH086654 (Nigg); R01MH115357 (Fair-Nigg); 2R37MH59105 (Nigg); R01 MH096773 (Fair); K23 MH108656 (Karalunas).

Data storage at OHSU is supported by the Oregon Clinical and Translational Research Institute funded by a grant from the National Center for Advancing Translational Sciences (NCATS), National Institutes of Health, through Grant Award Number UL1TR002369. The PDC (Perturbation of the Depression Connectome) research is supported by the National Institute of Mental Health (Grant No. U01MH110008). This manuscript reflects the views of the authors and may not reflect the opinions or views of the NIH, BANDA, PDC, HCP, or Oregon ADHD 1000 consortium investigators.

